# Physiological learning involves intrasynaptic reorganization of GluA1-containing AMPA receptors

**DOI:** 10.64898/2026.10.22.760928

**Authors:** Marijo Jevtić, Elena Hollergschwandtner, Bryan Kartono, Yui Sugiyama, Lenka Strnadová, Ragul Ramesh, Yugo Fukazawa, Makoto Itakura, Ryuichi Shigemoto

## Abstract

Learning and memory formation depend on persistent modifications of excitatory synaptic transmission in the hippocampus through changes in AMPA receptor (AMPAR) content and properties at potentiated synapses. However, directly demonstrating such changes after physiological, behavior-driven learning in the intact brain has remained technically difficult. Using SDS-digested freeze-fracture replica labeling (SDS-FRL), we examined the density and nanoscale localization of AMPAR subunits GluA1, GluA2, and GluA3 in the mouse hippocampal CA1 pyramidal cells. GluA2 density was higher than GluA1 and GluA3 densities at both synaptic and extrasynaptic sites, consistent with previous studies showing that AMPARs in CA1 pyramidal cells mostly contain GluA2. The extrasynaptic density of GluA3 was substantially lower than those of GluA1 and GluA2, pointing to distinct trafficking mechanisms. GluA1- and GluA2-containing receptors were enriched at synaptic peripheries, whereas GluA3 was more centrally localized and clustered, a subunit-specific intrasynaptic organization consistent with differential roles in plasticity. To investigate learning-induced reorganization, we combined the inhibitory avoidance, hippocampus-dependent contextual learning paradigm with SDS-FRL in the CA1 stratum radiatum. Learning induced a selective increase in synaptic GluA1 density accompanied by a concurrent decrease in GluA2 density, consistent with a compositional shift toward higher-conductance GluA1-containing receptors, specifically in the proximal CA1 area. Strikingly, we found a redistribution of GluA1 from the synaptic periphery toward the center, accompanied by increased clustering. These findings reveal intrasynaptic receptor redistribution alongside subunit composition change, a nanoscale molecular signature at CA3-CA1 synapses, which may constitute a mechanism of hippocampal memory formation in physiological learning.

**Significance Statement:** How the brain stores memories remains one of the most fundamental questions in biology. It is widely believed that memory formation strengthens specific synaptic connections in the hippocampus by increasing the number of postsynaptic AMPA receptors - the principal mediators of fast excitatory transmission. Here we show, using high-resolution electron microscopy, that physiological learning does not only increase GluA1 subunits but also selectively repositions GluA1-containing AMPA receptors within the synapse. Inhibitory avoidance training moved them toward the center with enhanced clustering specifically in the proximal CA1 region. This nanoscale reorganization induced by learning reveals a potential new dimension of synaptic memory storage in the brain.

## Introduction

The hippocampus became the primary focus of research to explain learning and memory formation after the case of patient H.M. (1), who was unable to form new long-term memory after the medial temporal lobectomy performed to treat epilepsy. The phenomenon of long-term potentiation (LTP), first described in the hippocampus (2), has since attracted enormous attention (3,4), based on the premise that LTP reveals a fundamental mechanism of memory encoding. Fast excitatory synaptic transmission underlying LTP is mediated by AMPA-type glutamate receptors (AMPARs), tetrameric ion channels assembled from GluA1–GluA4 subunits whose composition governs single-channel conductance, calcium permeability, trafficking, and interactions with postsynaptic scaffolding proteins (5). GluA1-containing receptors exhibit higher conductance and distinct activity-dependent trafficking compared to GluA2/3 heteromers (6,7), making subunit-specific redistribution a plausible and powerful mechanism for synaptic potentiation.

However, directly demonstrating that LTP is induced by physiological learning, and the resulting molecular changes at individual synapses, remains challenging. The difficulty lies in identifying the precise subregion and subpopulation of synapses affected, as effects may be subtle, sparse, and distributed across large areas surrounded by many unmodified synapses and therefore difficult to detect at the population level (8,9). The problem is compounded by the fact that learning can simultaneously induce LTP at some synapses and LTD at others, and by the inherent technical variability of available methods (8,9). Many hippocampus-dependent tasks also require multiple training trials, and differences in learning rates between animals can further obscure subtle synaptic changes. The inhibitory avoidance (IA) task offers practical solutions to several of these problems: IA training forms a stable memory trace in a single trial and produces robust changes in field excitatory postsynaptic potential, phosphorylation and delivery of GluA1, gene expression, and proteome in the dorsal CA1 area, indicating substantial and spatially localized synaptic plasticity (8,10–12).

The dorsal hippocampus is a key structure for acquiring and storing contextual fear memories (8,13), and these processes have been specifically linked to AMPAR trafficking and synaptic potentiation *in vivo* (8,14,15). Although functional differences along the proximodistal axis of CA1 are well established (16–20), learning-induced plasticity along this axis was previously unknown. Mitsushima and colleagues (21,22) demonstrated that AMPAR currents increase after IA training most prominently in CA1 proximal to CA2, with a weaker effect in distal CA1 in the rat. The observed increase in the AMPA/NMDA ratio — driven exclusively by AMPAR currents — can arise from several non-mutually exclusive mechanisms: increased receptor number at potentiated synapses, altered subunit composition, or more precise nanoscale alignment of AMPARs with the presynaptic neurotransmitter release machinery (23). Recent evidence has proposed that the PSD is organized into discrete nanodomains of 50–80 nm that are spatially aligned with presynaptic RIM/Munc13 scaffolds in trans-synaptic nanocolumns, and that LTP could drive AMPARs into closer apposition with release sites — a nanoscale redistribution that enhances transmission independently of changes in total receptor number (24–28).

In this study, we investigated the detailed distribution of AMPAR subunits in mouse CA1 pyramidal cells, with special emphasis on CA3-CA1 synapses, utilizing high-specificity antibodies against GluA1, GluA2, and GluA3 combined with SDS-digested freeze-fracture replica labeling (SDS-FRL) — the highest-resolution quantitative approach currently available for *in situ* visualization of membrane proteins (29) — and our in-house image analysis platform DAREA (30). We first establish the baseline nanoscale organization of all three subunits across the radial axis of CA1 and within individual synapses, revealing subunit-specific differences in intrasynaptic positioning. We then combine IA training with SDS-FRL to show that physiological learning selectively increases synaptic GluA1 density and drives its redistribution from the synaptic periphery toward the center with enhanced clustering, specifically in the proximal CA1 stratum radiatum. These findings implicate intrasynaptic receptor reorganization alongside changes in subunit composition, as a mechanism underlying hippocampal memory formation.

## MATERIALS AND METHODS

### Animals

For this study, wild-type (WT) male C57BL/6J mice aged 8–10 weeks were used. Before experiments, animals were kept on a 12:12 light-dark cycle with ad libitum access to food and water in the Preclinical Facility of IST Austria. All experiments were performed in accordance with the license approved by the Austrian Federal Ministry of Science and Research (Animal license number: BMWFW-66.018/0012-WF/V/3b/2016) and the Austrian and EU animal laws.

### Labeling reagents

Antibodies used in this study are summarized in Table 1.

**Table 1.** Labeling reagents used in this study:

| Reagent | Concentration | Reference/Source |
| --- | --- | --- |
| Anti GluA1 rabbit antibody | 4 µg/ml | Antal et al, 2008 (31) |
| Anti-GluA2 rabbit antibody | 2 µg/ml | Tabata et al, 2019 (32) |
| Anti-GluA3 rabbit antibody | 10 µg/ml | Rubio et al, 2017(33) |
| Anti-panAMPA rabbit antibody | 4 µg/ml | Eguchi et al, 2020 (34) |
| Anti-rabbit antibody, 5 nm gold conjugated | 1:30 dilution | British Biocell International |

### SDS-digested freeze-fracture replica labeling (SDS-FRL)

SDS-FRL technique was described previously (35). Animals were anesthetized with a lethal dose of pentobarbital (200 mg/kg), and brains were fixed via transcardial perfusion with either 2% (Figure 1) or 0.5% paraformaldehyde (PFA) (all other figures), 15% picric acid in 0.1 M phosphate buffer (PB). Mouse brains were sliced into coronal sections at a thickness of 80 µm (Figure 1) or 120 µm (Figures 2–4) using vibratome (Linear Slicer Pro, Dosaka). Brain slices were cryoprotected with 30% glycerol in 0.1 M PB. First, the CA1 area from the dorsal hippocampus was trimmed out, and then high-pressure frozen using the Baltec HPM010. Frozen tissue samples were fractured using the Baltec BAF060 and/or Jeol JFDV and a thin layer of carbon (5nm) was deposited on top of fractured tissue, followed by 2 nm platinum layer and additional 20nm of carbon. Tissue was then dissolved in SDS solution (2.5% sodium dodecyl sulfate solution (SDS), 20% sucrose, 0.05% sodium azide in 15 mM Tris buffer) in a shaking incubator at 80°C and 30-50 RPM. For labeling, replicas were washed once in SDS solution, then once in washing buffer (WB, 0.1% Tween-20, 0.05% BSA (bovine serum albumin), 0.05% NaN_3_ in TBS (Tris-buffered saline), twice in TBS and then incubated in blocking buffer (BB, WB with 5% BSA) for 30-60 min. After blocking, replicas were incubated in primary antibody solution (GluA1, GluA2, GluA3, or panAMPA (specific to GluA1-3) antibody, diluted in BB) for 24-72h at 15°C. The specificity of the antibodies for GluA1, GluA2, and GluA3 has been confirmed using respective knockout (KO) animals (GluA1 (31), GluA2 (32), GluA3 (33)). After washing in 1x WB and 2x TBS, replicas were incubated in BB and then, in secondary antibody solution (gold conjugated secondary antibody diluted in BB, see Table 1). After washing in WB, 2x TBS, replicas were flattened on the surface of MilliQ water before mounting them on copper grids with parallel bars for EM observation.

### Modified SDS-FRL protocol for intrasynaptic analysis

For the quantification of AMPAR numbers and analysis of intrasynaptic distribution of AMPAR subunits, we prepared an additional set of samples using a slightly modified “flattening” protocol (29). This method allows for efficient sampling of spine synapses with complete PSD areas, which are necessary for the analysis of center-periphery preference. Labeling was performed in the same way as above using 0.5% PFA for perfusion and thicker slices (120 µm) for the high-pressure freezing, resulting in a slight flattening of slices before freezing. Synapses with complete PSDs and at least 4 gold particles were sampled from the middle one-third of the stratum radiatum. Although higher density values of observed labeling were likely due to weaker fixation of samples and/or the omission of synapses with less than 4 gold particles, the relative density between different subunits was consistent.

### Grid-glued replica

Samples for GluA1 labeling in the IA task were prepared using modified, “glued replica” protocol (36) in order to preserve complete CA1 areas for analysis, albeit at the cost of reduced labeling density. Only replicas containing intact proximal and distal parts of the CA1 area were sampled and analyzed. After fracturing, the replicated side of the samples (tissue + replica) were glued to nickel finder grids with a UV (ultraviolet)-activated glue (optical adhesive 61, Norland, Cranbury, NJ). Tissue digestion and replica labeling were done as described above. In the case of the grid-glued replica, it is necessary to dissolve glue before EM observation. To protect gold particle labeling, samples were coated with a 20nm carbon layer before treating the samples with a glue-dissolving agent (Dynasolve 711, Dynaloy, Indianapolis, IN) for 2x 45 min at 60°C and 50 RPM in a shaking incubator.

### EM imaging

Postsynaptic sites on the E-face of replicas were identified as tightly packed clusters of intramembrane particles (IMPs), representing postsynaptic membrane specializations (PSDs). The vast majority of synapses found on dendritic spines in the stratum radiatum of the CA1 area have their presynaptic origin in the CA3 area, while synapses found on dendritic spines in the stratum lacunosum-moleculare have their origin mainly from the entorhinal cortex (37,38).

Images from extrasynaptic sites in the pyramidal cell layer were taken from the E-face of pyramidal cell somata, each image representing a different cell. In all other layers, extrasynaptic images were sampled from the E-face of dendritic shafts after confirming that the observed dendrite belongs to the CA1 pyramidal cells (by presence of spine necks).

For Figure 1, only replicas containing all four strata, the stratum oriens (**ORI**), stratum pyramidale (**PCL**), stratum radiatum (SR), and stratum lacunosum-moleculare (**LM**), were selected for analysis. Predefined subregions within the stratum pyramidale were treated as individual sampling units. Within the stratum radiatum, the proximal radiatum (**PR**) was defined as the portion closer to the stratum pyramidale, whereas the distal radiatum (**DR**) was defined as the portion closer to the stratum lacunosum-moleculare. Pyramidal cell bodies, dendrites, and synapses were randomly sampled within each designated anatomical unit. From each designated sampling unit, 20–30 images of synaptic sites, including postsynaptic density (PSD) areas, and 10–15 images of extrasynaptic sites, including cell bodies and dendrites, were collected and analyzed. Each replica represented an individual animal, and a total of four animals were analyzed.

For Figure 2, synapses were randomly sampled from the middle third of the stratum radiatum (middle radiatum). A total of 15–25 synapses were sampled per replica. Each replica represented one animal, and four animals were analyzed.

For Figures 3–4, only replicas containing intact proximal and distal portions of the CA1 area were included in the analysis. Synapses were randomly sampled from the middle third of the stratum radiatum in CA1 regions proximal and distal to the CA2 area. For each region of interest within the same replica (proximal CA1 and distal CA1), 20–30 synapses were sampled. Each replica represented an individual animal, and 4–8 animals were analyzed per dataset. Group sizes were six animals for GluA1-trained, eight animals for GluA1-untrained, and four animals per group for the trained and untrained GluA2, GluA3, and panAMPA labeling datasets.

Image acquisition and sampling were performed by an investigator blinded to the experimental group and sample identity. Images were acquired using Tecnai 10 or Tecnai 12 (FEI) transmission electron microscopes.

### Image analysis, visualization and statistics

For image analysis and visualization GPDQ (39) or Darea (30) software was used. First, areas of interest (PSD) were demarcated and gold particles were detected manually in the replica images. Subsequently, gold particle number and density within the demarcated area and the outer rim of synapses, distances between particles, and the center of gravity for each demarcated area were analyzed. For each gold particle within a demarcated area, the center periphery index (CPI) was determined using the DAREA software, and the mean was calculated for each synapse. Center periphery index is calculated using a formula: 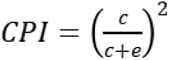 where *c* indicates distance from the gold particle to the center of gravity of the synapse, while *e* indicates distance to the nearest edge of the demarcated synapse. Clustering analysis was performed using DBSCAN (minPts = 3) on receptor coordinates obtained from single-synapse datasets. The neighborhood radius (ε) was determined empirically using k-distance analysis (k = minPts) across all synapses (40). The distance from each particle to its k-th nearest neighbor was calculated, and the resulting distances were sorted in ascending order to produce a k-distance plot. The value of ε was selected at the point of maximum curvature (the “elbow”) of the plot, representing the transition between dense clusters and sparse observations. This ε value was then used as the neighborhood radius for the DBSCAN analysis. In parallel, nearest-neighbor distance (NND) distributions were also computed, yielding a median NND of ∼17–40 nm. Based on these analyses, ε values in the range of 25–60 nm (depending on the labeling density, approximately 1.5–2× median NND) were selected to capture full extent of each receptor cluster while avoiding over-fragmentation at smaller ε and artificial merging at larger ε. Clustering results were verified to be robust across this ε range. Final ε chosen was 40 nm for Figure 2 and 50 nm for Figure 4.

For each synapse, clustered particles were defined as those assigned to DBSCAN clusters, and the fraction of clustered particles was calculated as the ratio of clustered to total particles. Cluster size was quantified as the number of particles per cluster, considering only clusters with ≥3 particles. Cluster area was estimated as the convex hull area of particles for each cluster and normalized to the total synapse area to account for differences in synapse size. Nearest-neighbor distances were additionally normalized by local particle density (multiplying by the square root of density) to enable comparison across synapses with varying densities. All metrics were first computed per synapse and subsequently averaged per animal for statistical analysis. Significance of gold particle clustering was analyzed by comparing real distributions with random gold particle distributions obtained by Monte Carlo simulation. We used the same shape, size and number of particles as a real synapse data to run the simulation 100 times. Average of these 100 repeated simulations were used to compare with the real experimental data for each synapse.

Statistical analysis of data was performed by GraphPad Prism 6 (San Diego, CA, USA). Graphs were also prepared with GraphPad Prism. Error bars indicate standard error of the mean. The p-values, * *p <* 0.05, ** *p <* 0.01, and *** *p <* 0.001 were indicated in the graphs. Detailed statistics can be found in the Supplementary Information.

### Inhibitory avoidance task

The inhibitory avoidance (IA) task was designed as previously described (21,22). Animals were placed in the light chamber of the IA apparatus for habituation for 5 minutes one day before the experiment. On the day of the experiment, animal was placed in the light compartment of the apparatus and 5s later the door to the dark compartment was open so the animal could freely pass. Time from opening the door until the animal entered the dark compartment was measured as latency **before** the electric shock. Upon entering the dark compartment, 3s later the door was closed and electric foot shock (1s, 0.5mA) was applied via electrified steel rods in the floor of the apparatus. Animal was kept in the dark compartment for 10s and then returned to the home cage. After 30 min, the animal was again placed in the light compartment of the apparatus. The time to enter the dark compartment was measured as latency **after** the shock and assessed as a measure of successful learning. For electron microscopy analysis, animals were anesthetized and perfused shortly after the memory retrieval, and fixed brains from animals with the latency of at least 100 seconds during second exposure (after shock) were selected for further processing. Control (untrained) animals were kept in their home cages before perfusion, without exposure to the IA task.

### Use of Generative AI

ChatGPT (GPT-5.2) and Claude (Sonnet 4.6) were used during initial manuscript preparation to generate a preliminary draft and organize the text’s conceptual structure. The authors subsequently revised, rewrote, and critically edited the AI-assisted material, ensuring scientific accuracy and appropriate interpretation. The authors take full responsibility for the accuracy and integrity of the final manuscript.

## RESULTS

### Extrasynaptic and synaptic distribution of AMPAR subunits along the radial axis in CA1

To determine the spatial distribution of AMPAR subunits across the CA1 radial axis, we utilized a quantitative EM technique (SDS-FRL) (29,30,35), which allows for *in situ* visualization of gold-particle-immunolabeled membrane proteins, combined with advanced image analysis tool developed in-house (DAREA) (30) for subsequent analysis of gold particle distribution parameters. We quantified gold particles for GluA1, GluA2, and GluA3 in defined laminar compartments, which include the stratum oriens (ORI), pyramidal cell layer (PCL), proximal (closest to PCL), middle, and distal stratum radiatum (PR, MR, DR), and stratum lacunosum-moleculare (LM) (Figure 1A). Two-way ANOVA of sampled PSD area revealed significant difference between layers (F(4, 60) = 5.677, *p* < 0.001, n = 4 animals per group), no difference between subunits (F(12,60) = 0.502, *p* = 0.91) and no interaction (F(3,60) = 0.1570, *p* = 0.92), indicating representative sampling of the PSD across subunits. The post hoc analysis indicated that the average PSD area was largest in the stratum lacunosum-moleculare, which was common across all types of labeling (Supplementary Figure S3A and Supplementary table S1) and is consistent with previous studies (37,41).

Qualitative inspection of representative electron micrographs confirmed the presence of immunogold labeling at both synaptic and extrasynaptic membranes across all layers (Figure 1B-D). Based on the previous results showing that AMPA receptors expressed in CA1 pyramidal cells are Ca^2+^-impermeable GluA2-containing receptors (42,43), we first examined the labeling efficiency of the GluA2 antibody relative to the panAMPA antibody, which has close to 100% labeling efficiency for functional AMPAR channels (34). The distribution of gold particles for panAMPA showed similar patterns as those for GluA2 in synaptic and extrasynaptic sites across all layers (Supplementary Figure S1A-D). When densities of panAMPA and GluA2 particles were compared in replicas prepared from the same animals to control for technical variability, no significant difference was detected in synaptic densities (Supplementary Figure S1E, panAMPA vs GluA2, *p* = 0.13, unpaired t-test). This result indicates that the labeling efficiency of the two antibodies are similar, assuming that only one antibody can bind to one AMPA receptor channel. To examine the labeling efficiencies of the GluA1 and GluA3 antibodies, we next compared the particle densities obtained with the GluA1 and panAMPA antibodies in complementary replicas prepared from GluA3 knockout (KO) mice, and those with the GluA3 and panAMPA antibodies in complementary replicas prepared from GluA1 KO mice, assuming that AMPA receptors of only GluA1-GluA2 or GluA3-GluA2 subunit compositions are expressed in CA1 pyramidal cells in GluA3 or GluA1 KO mice, respectively. We found no significant differences in the labeling density between these antibodies (Supplementary Figure S1F-G, panAMPA vs GluA1 in GluA3KO, *p* = 0.34, n = 32 synapses per group, panAMPA vs GluA3 in GluA1KO, *p* = 0.21, n =26 synapses for panAMPA, n = 20 synapses for GluA3, unpaired t-test). This situation allowed us to compare the subunit densities across GluA1-3 subunits.

Based on the similar labeling efficiencies of our GluA1, GluA2, and GluA3 antibodies, we compared absolute densities of gold particles for these subunits in synaptic and extrasynaptic (cell bodies and dendrites) membranes (Table 2). Two-way ANOVA of GluA densities in synapses revealed significant main effects of subunit (F(2,45) = 13.48, *p* < 0.0001) and layer (F(4,45) = 2.696, *p* = 0.04, n = 4 animals per group), with no significant subunit × layer interaction (F(8,45) = 0.1230, p = 0.998). Post hoc Tukey’s multiple-comparison tests revealed that synaptic GluA2 density was significantly higher than both GluA1 (*p* < 0.001) and GluA3 (*p* < 0.001), whereas no significant difference was observed between GluA1 and GluA3 (*p* = 0.989). Similarly, two-way ANOVA of extrasynaptic densities revealed significant main effects of subunit (F(2,54) = 27.04, p < 0.0001) and layer (F(5,54) = 2.753, *p* = 0.03, n = 4 per group), with no significant subunit × layer interaction (F(10,54) = 0.5110, *p* = 0.88). Post hoc Tukey’s multiple-comparison tests revealed that extrasynaptic GluA2 density was significantly higher than both GluA1 (*p* < 0.001) and GluA3 (p < 0.001), and that GluA1 density was significantly higher than GluA3 (*p* = 0.03).

To compare relative GluA densities in synapses across layers, we normalized the density values to that in the stratum oriens (Figure 1E-G). The normalized synaptic GluA densities were consistently higher throughout the stratum radiatum than in the flanking layers, stratum oriens and stratum lacunosum-moleculare. In contrast, normalized extrasynaptic densities (Figure 1F) showed a layer-dependent gradient with increasing densities from the pyramidal cell layer through the stratum radiatum, while stratum oriens and stratum lacunosum-moleculare displayed similar intermediate densities.

Furthermore, the ratios of extrasynaptic and synaptic densities for each GluA subunit across layers indicated consistently lower extrasynaptic pool of GluA3-containing receptors (Figure 1G). Two-way ANOVA revealed significant main effects of subunit (F(2,45) = 15.05, *p* < 0.001) and layer (F(4,45) = 3.93, *p* = 0.008, post hoc comparison in Supplementary Table S2) with no interaction (F(8,45) = 0.30, *p* = 0.96, n = 4 animals per group). This result indicates that the GluA1-GluA2 receptor occupies the major population of extrasynaptic AMPARs. To examine the contribution of GluA1 to extrasynaptic AMPARs, we compared the extrasynaptic/synaptic ratios of panAMPA receptor labeling between wild-type (WT) and GluA1 KO mice in the middle stratum radiatum (Supplementary Figure S2). We found significantly lower ratio in GluA1 KO than WT mice (Supplementary Figure S2B, *p* = 0.02, n = 5 animals for WT and 3 for KO, unpaired t-test), indicating that GluA1 is necessary for keeping the high ratios of extrasynaptic AMPARs. In addition, the reduced ratio for panAMPA labeling in GluA1 KO mice was similar to that for GluA3 labeling in WT mice (Supplementary Figure S2B), indicating that the allocation of GluA2-GluA3 receptors in the extrasynaptic sites remained low in the absence of GluA1.

**Table 2.** Absolute values for AMPAR subunit labeling in individual CA1 strata in synaptic and extrasynaptic sites.

| Synaptic | s.oriens | s.pyramidale | s.radiatum (proximal) | s.radiatum (middle) | s.radiatum (distal) | s.lacunosum-moleculare |
| --- | --- | --- | --- | --- | --- | --- |
| GluA1 | 334.9±67.7 | N/A | 462.4±64.9 | 443.3±53.7 | 508.3±75.8 | 314.1±60.9 |
| GluA2 | 616.1±101.5 |  | 881.2±204.5 | 787.4±166.6 | 831.5±184.9 | 532.3±99.7 |
| GluA3 | 333.3±65.4 |  | 492.1±149.7 | 426.7±54.9 | 479.6±119.7 | 279.9±59.8 |
| Extrasynaptic | s.oriens | s.pyramidale | s.radiatum (proximal) | s.radiatum (middle) | s.radiatum (distal) | s.lacunosum-moleculare |
| GluA1 | 54.6±15.3 | 34.7±6.5 | 46±13.3 | 64.6±14.2 | 69.7±17 | 51±10.5 |
| GluA2 | 97.2±17.1 | 60.5±13.2 | 85.7±19.6 | 131.1±30.7 | 118.3±27.5 | 76.4±14.2 |
| GluA3 | 36.7±9.2 | 20.4±7.8 | 25.3±10 | 34.6±8.7 | 36.3±8.6 | 27.7±8.5 |

**Figure 1.**
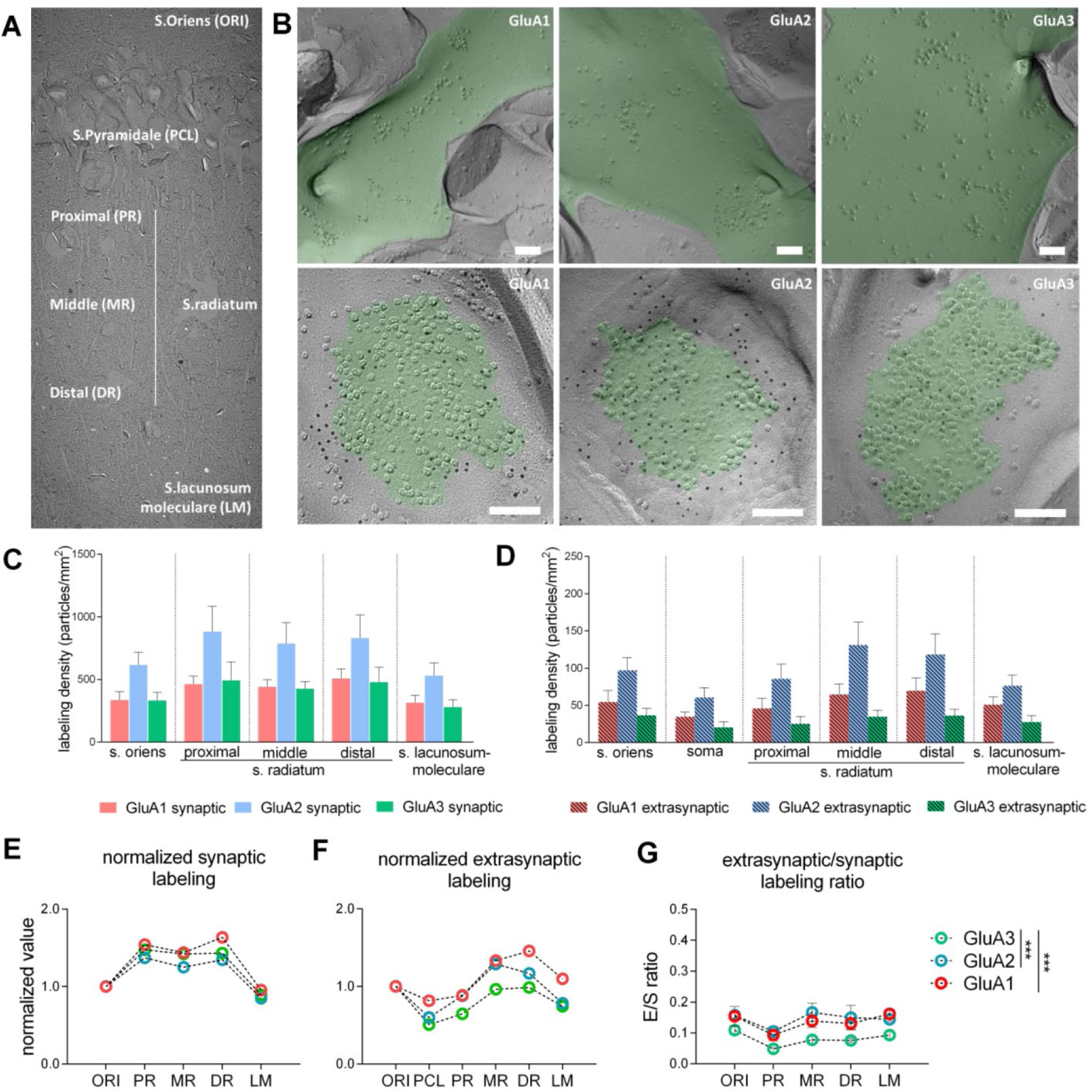
Radial distribution and synaptic versus extrasynaptic localization of AMPAR subunits in CA1. **A.** Low magnification replica image of the mouse CA1 area of the hippocampus, including stratum oriens (ORI), pyramidal cell layer (PCL), proximal, middle, and distal stratum radiatum (PR, MR, DR), and stratum lacunosum-moleculare (LM). **B.** Representative electron micrographs illustrating immunogold labeling for GluA1, GluA2, and GluA3 in synaptic (bottom) and extrasynaptic (top) sites. **C-D.** Quantification of gold particle densities (particles/µm²) for synaptic (left) and extrasynaptic (right) compartments across layers. Data are presented as mean ± SEM. Absolute values are directly comparable across subunits based on similar labeling efficiency of antibodies for GluA1-3. **E-F.** Left and middle panels show synaptic and extrasynaptic densities normalized to that in the stratum oriens for each sample, plotted across layers. **G.** The ratios of the extrasynaptic to synaptic densities of GluA1-3. Data are presented as mean ± SEM (n = 4). Two-way ANOVA revealed significant effects of both layer and subunit, with no interaction. Post hoc comparisons revealed significantly lower ratios for GluA3 than GluA1 and GluA2. Asterisks indicate levels of statistical significance: *p* < 0.001 (***).

### Distinct intrasynaptic distribution of AMPAR subunits

To investigate patterns of gold particle distribution within postsynaptic sites, we prepared a separate dataset of synapse images exclusively from the stratum radiatum and utilized DAREA software to perform a series of analyses including simulation of random distribution of gold particles using the same PSD area and number of gold particles detected in the real dataset, as indicated in flowchart in the Figure 2A.

Consistent with the results shown in Figure 1, we found higher labeling density for GluA2 (1126 ± 54.5 particles/µm²) than for both GluA1 (636.7 ± 41.7 particles/µm²) and GluA3 (613.2 ± 35.1 particles/µm²) (Figure 2B; one-way ANOVA, F(2,9) = 7.75, *p* = 0.01, n = 4 animals per group, post hoc Tukey’s multiple comparisons test, p = 0.02 for GluA1 vs GluA2, *p* = 0.02 for GluA2 vs GluA3). To ensure that differences in labeling density were not attributable to sampling bias, we confirmed that synaptic areas did not differ significantly between GluA1, GluA2, and GluA3 samples (Supplementary Figure S3B, one-way ANOVA, F(2,9) = 2.23, *p* = 0.16, n = 4 animals per group).

To quantify radial positioning of AMPAR subunits within individual synapses, we used a center-periphery index (CPI) (30), which indicates the particle position relative to synaptic center (CPI = 0) and the edge (CPI = 1). Mean CPI values differed significantly between subunits (Figure 2C; one-way ANOVA, *F*(2,9) = 9.58, *p* = 0.006, n = 4 animals per group), indicating subunit-specific radial organization. Post hoc analysis (Tukey’s multiple comparisons test) revealed higher CPI values for GluA1 (0.68 ± 0.01, *p* = 0.001) and GluA2 (0.67 ± 0.01, *p* = 0.01) than for GluA3 (0.52 ± 0.01), whereas GluA1 and GluA2 did not differ. Mean CPI values for GluA1 and GluA2, but not GluA3, were significantly higher than those of density-matched simulated random distributions (Supplementary Figure S4A: GluA1, *p* = 0.01, n = 4 animals; GluA2, *p* = 0.02, n = 4 animals; GluA3, *p* = 0.3, n= 4 animals per group, paired *t*-tests).

Cumulative CPI distributions also differed between subunits, with GluA3 shifted toward lower CPI values compared with both GluA1 and GluA2, whereas GluA1 and GluA2 did not differ (Figure 2D: GluA1 vs GluA3, *D* = 0.469, *p* < 0.001, n = 87 vs 88 synapses, respectively; GluA2 vs GluA3, *D* = 0.433, *p* < 0.001, n= 81 vs 83, respectively; GluA1 vs GluA2, *D* = 0.080, *p* = 0.92, n = 87 vs 81 synapses, respectively, Kolmogorov–Smirnov tests). Relative to simulated random distributions, GluA1 and GluA2 were shifted toward higher CPI values, whereas GluA3 was shifted toward lower CPI values (Supplementary Figure S4B: GluA1 vs simulated, *D* = 0.410, *p* < 0.001, n = 87 synapses; GluA2 vs simulated, *D* = 0.436, *p* < 0.001, n = 81 synapses; GluA3 vs simulated, *D* = 0.477, *p* < 0.001, n = 88 synapses, Kolmogorov–Smirnov tests). These results indicate preferential localization of GluA1 and GluA2 toward the synaptic periphery and of GluA3 toward the synaptic center.

Furthermore, the proportion of particles within the 30 nm perisynaptic outer rim differed significantly between subunits (Figure 2E; *F*(2,9) = 7.80, *p* = 0.01, n = 4 animals per group, one-way ANOVA). Post hoc analysis (Tukey’s multiple comparisons test) revealed a higher proportion of GluA1 (18.42 ± 1.59%, *p* = 0.02) and GluA2 (17.89 ± 2.62%, *p* = 0.02) particles in the outer rim than GluA3 (8.51 ± 1.58%), whereas GluA1 and GluA2 did not differ, further confirming preferential localization of GluA1 and GluA2 toward the synaptic periphery and of GluA3 toward the synaptic interior.

### Subunit-specific differences in receptor clustering

Nearest-neighbor distance (NND) analysis was used to assess clustering properties of GluA subunits (44). Compared with density-matched simulated random distributions, GluA3 particles exhibited significantly shorter NND values (Figure 2F; *p* = 0.009, n = 4 animals, paired t-test), indicating local clustering of GluA3-containing AMPARs. GluA1 showed a tendency toward shorter NND values (*p* = 0.16, n = 4 animals), whereas GluA2 did not differ from its corresponding simulated random distribution (*p* = 0.46, n = 4 animals).

Clustering was further quantified with DBSCAN analysis implemented in DAREA software. The GluA3 distribution exhibited a significantly higher proportion of clustered particles than the simulated distribution (Figure 2G, *p* = 0.003, n = 4 animals, paired t-test, whereas GluA1 and GluA2 showed non-significant trends (*p* = 0.10 and *p* = 0.12, respectively, n = 4 animals per group). Analysis of particle number per cluster further revealed subunit-specific differences. GluA3 exhibited significantly higher number per cluster compared to simulated random distributions (Figure 2H, *p* = 0.04, n = 4 animals, paired t-test), whereas GluA1 showed a non-significant trend (*p* = 0.17, n = 4 animals) and GluA2 did not differ from random (*p* = 0.88, n = 4 animals). We also analyzed the center-periphery placement of DBSCAN-defined cluster centroids. Comparison with simulated distributions showed that GluA1 and GluA2 clusters were positioned more peripherally than expected by random (*p* < 0.001 and *p* = 0.02 for GluA1 and GluA2, respectively, n = 4 animals per group, paired t-test), whereas GluA3 cluster centroids did not differ from random (*p* = 0.40, n = 4 animals) (Figure 2I).

Together, these analyses demonstrate that AMPAR subunits exhibit distinct nanoscale organization within synapses. GluA3 is characterized by a compact, centrally localized organization with significant local clustering. In contrast, GluA1 is preferentially localized toward the synaptic periphery and enriched within the outer synaptic rim. GluA1 showed a tendency toward local clustering while GluA2 exhibited more diffuse spatial distribution.

**Figure 2.**
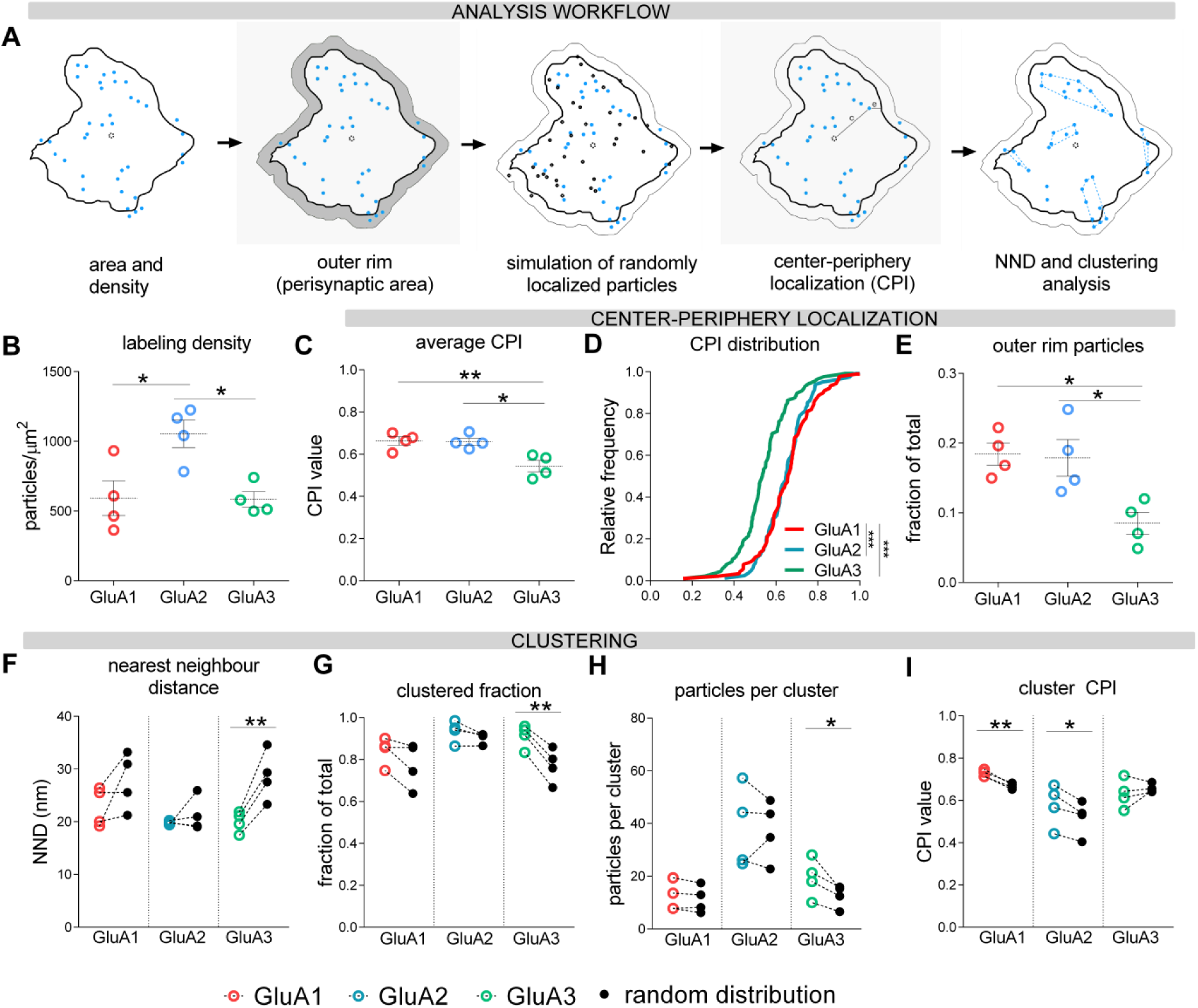
Subunit-specific nanoscale organization of AMPARs within CA1 synapses. **A.** Schematic overview of analysis workflow, including identification of synaptic profiles, particle detection, generation of simulated random distributions, calculation of center–periphery index (CPI), nearest-neighbor distance (NND), and clustering analysis using DBSCAN. Representative demarcation of synaptic area includes gold particles (blue dots), outer rim (gray area), simulated randomly distributed gold particles (black dots) and detected gold particle clusters (dotted blue line). **B.** Labeling density across subunits showing higher density of GluA2 than GluA1 and GluA3. **C.** Center–periphery index (CPI) for GluA1, GluA2, and GluA3. Individual data points represent biological replicates; lines indicate mean ± SEM. **D.** Cumulative distribution functions (CDFs) of CPI values, showing overlapping distributions for GluA1 and GluA2 and a distinct distribution for GluA3. **E.** Fraction of particles located within the perisynaptic outer rim, indicating lower perisynaptic density of GluA3 than GluA1 and GluA2. **F.** Real NND values compared to those of simulated random distributions. Paired points represent matched real and simulated values. GluA3 shows significantly lower NNDs than simulated NNDs, indicating clustering property. **G.** Fraction of clustered particles relative to simulated values, indicating significant clustering of GluA3. **H.** Number of particles per cluster, indicating differences in clustering strength across subunits. **I.** CPIs of cluster centroids showing peripheral localization for GluA1 and GluA2 clusters. All summary plots show mean ± SEM. Statistical significance was assessed using one-way ANOVA with post hoc comparisons, paired t-tests for comparisons to simulated distributions, and Kolmogorov–Smirnov tests for distribution analyses. Asterisks indicate levels of statistical significance: p < 0.05 (*), p < 0.01 (**), p < 0.001 (***).

### Changes in subunit density induced by inhibitory avoidance task

To determine whether learning induces region-specific changes in synaptic AMPAR organization in the CA1 area, mice were subjected to an inhibitory avoidance (IA) task and analyzed 30 min after the retrieval (Figure 3A). Using the behavioral procedure for rats described in Mitsushima et al. (21,22) with some modifications, we performed IA training using male WT C57BL/6 mice. The trained animals exhibited a significant increase in time spent in the light compartment following the foot shock (Figure 3B: p < 0.001, n = 19 animals, paired t-test), demonstrating the robust and stable memory formation, which persisted for at least 72h. (Supplementary Figure S5).

Based on the reported regional differences in the rat (21), we investigated the synaptic labeling density for GluA subunits in the proximal (adjacent to CA2) and distal (adjacent to subiculum) CA1 regions of IA trained and untrained animals (Figure 3C). Immunogold labeling was quantified in the middle one-third of the stratum radiatum in these regions within the same replica samples to control for technical variability. In the IA trained animals, the labeling density for GluA1 was markedly higher in the synapses in the proximal compared to the distal CA1 (Figure 3D: 239± 27.2 particles/µm^2^ in proximal vs 111±14.5 particles/µm^2^ in distal regions, *p* = 0.001, n = 6 animals, paired t-test), whereas that for GluA2 was slightly but significantly lower (Figure 3F, 436.7±115.4 particles/µm^2^ in proximal vs 531.6±136.4 in distal regions, *p* = 0.02, n = 4 animals). No significant changes in the labeling density for GluA3 and panAMPA was detected (GluA3: Figure 3H, 280.4±54.38 in proximal and 254.3±26.09 in distal regions, *p* = 0.71, n = 4 animals; panAMPA: Supplementary Figure S6B, 744.2±157.2 in proximal and 796.3±198.0 in distal regions, *p* = 0.75, n = 4 animals, paired t-test). In control animals without IA training, labeling for all three subunits was comparable between the proximal and distal CA1 regions (Supplementary Figure S6A).

To facilitate direct comparison between subunits, we calculated ratios of particle density values in the proximal and distal CA1 regions (P/D ratio). The P/D ratio was significantly higher in the trained than control animals for GluA1 (Figure 3E, p < 0.001, n = 6 animals per group, unpaired t-test), whereas that for GluA2 showed a lower trend in the trained than control animals (Figure 3G, *p* = 0.09, n = 4 animals per group). The P/D ratio for GluA3 exhibited no significant difference between the trained and untrained animals (Figure 3I, *p* = 0.71, n = 4 animals per group).

Although the P/D ratio for panAMPA showed no significant difference between the trained and untrained animals (Supplementary Figure S6B, *p* = 0.81, n = 4 animals per group), it showed a trend of higher value than that for GluA2 (Supplementary Figure S6C, *p* = 0.08, n = 4 animals per group, unpaired t-test), which could imply an increase of GluA2-lacking AMPARs in the proximal CA1 region. The proximal/distal ratios of PSD areas were not significantly different between the groups for any GluA subunit (Figure 3E, 3G, 3I), indicating that the density differences described above do not simply reflect changes in synapse size.

These findings indicate that the IA training induces subunit-specific proximodistal differences of synaptic AMPAR expression in the CA1 region of the hippocampus.

**Figure 3.**
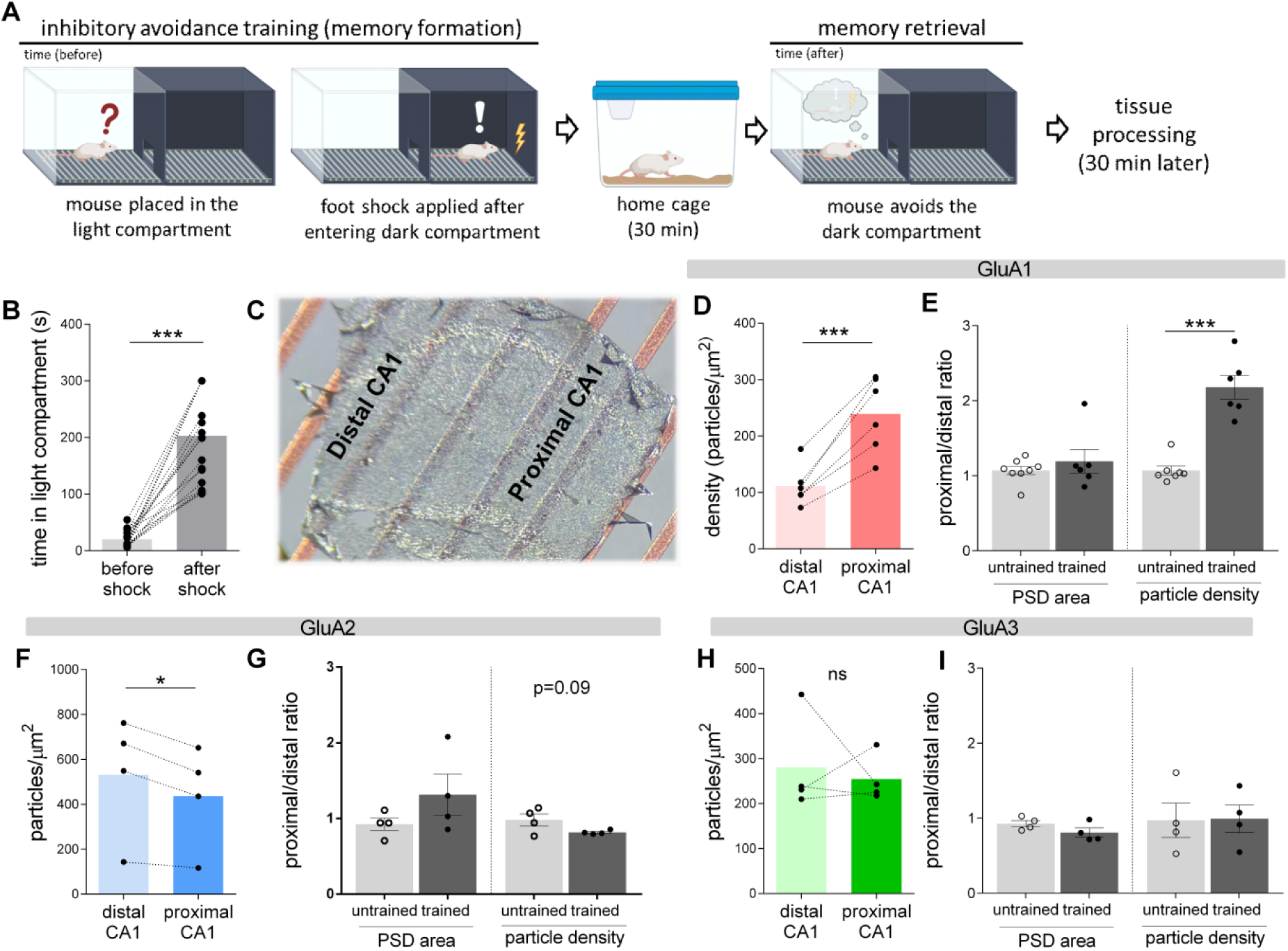
Learning induces proximodistal asymmetry of GluA1 in CA1. **A.** Schematic of the inhibitory avoidance training and memory retrieval procedure. Mice were placed in a light compartment and allowed to enter a dark compartment where a mild foot shock was delivered. Animals were tested 30 min after training. **B.** Behavioral performance during inhibitory avoidance. Time spent in the light compartment before and after the electrical shock is shown for individual animals (paired measurements). Lines connect values from the same animal. Statistical significance was assessed using a paired t-test. **C.** Schematic representation of proximal and distal CA1 regions analyzed. Proximal CA1 is defined as the region adjacent to CA2, whereas distal CA1 is located toward the subiculum. **D-I.** Proximodistal differences of GluA1-3 labeling densities in CA1 (panels D, F, H). Data are shown as paired measurements from the same animals for trained groups. Lines connect proximal and distal values within each animal. Ratio of proximal/distal values is shown in panels E (GluA1), G(GluA2) and H (GluA3). Statistical significance was assessed using paired t-tests (proximal vs distal) and unpaired t-tests (untrained vs trained). All data are presented as individual data points with mean ± SEM. Proximal and distal measurements were obtained from the same replicas, providing internal normalization for each animal.

### Changes in intrasynaptic organization after inhibitory avoidance task

To determine how learning reorganize the nanoscale distribution of GluA1-containing AMPA receptors within synapses, we analyzed their radial distribution and clustering properties in the middle stratum radiatum of proximal and distal CA1 regions of the IA trained animals. Analysis of radial localization revealed a consistent redistribution of GluA1 from the synaptic periphery toward the center. Although the mean CPI values showed only a non-significant trend of reduction (Figure 4A, *p* = 0.14, n = 6 animals, paired t-test), cumulative distribution analysis of the CPI demonstrated a pronounced shift toward lower values in proximal compared with distal synapses (Figure 4B, D = 0.143, *p* < 0.001, n = 65 and 61 distal and proximal synapses, respectively, Kolmogorov–Smirnov test). Consistent with this, the fraction of particles located in the perisynaptic outer rim was also significantly reduced in proximal compared to distal synapses (Figure 4C, *p* = 0.004, n = 6 animals, paired t-test).

We next assessed clustering property of GluA1 using the NND analysis. In both distal and proximal synapses, real NND values were significantly lower than simulated random distributions in trained animals (Supplementary Figure S7: distal, *p* = 0.001, n = 6 animals; proximal, *p* = 0.02, n = 6 animals, paired t-test), indicating significant clustering of GluA1-containing receptors. As previously observed (45), density differences (Figure 3) can mask the true intrasynaptic spatial reorganization. Thus, we focused our NND and DBSCAN clustering analyses on density-corrected or density-independent metrics. Density-corrected NND distributions revealed a distinct pattern of distribution between proximal and distal synapses (Figure 4D: D = 0.275, *p* = 0.017, n = 65 and 61 distal and proximal synapses, respectively, Kolmogorov–Smirnov test), indicating a reorganization of receptor spatial distribution independent of receptor density. Notably, density-corrected cluster area distributions exhibited a shift toward larger values in proximal synapses (Figure 4E: D = 0.476, *p* < 0.001), demonstrating an expansion of clustered domains. Additionally, the center-periphery index of clusters was significantly lower in proximal compared to distal synapses (Figure 4F: *p* = 0.03, n = 65 and 61 distal and proximal synapses, respectively, paired t-test), demonstrating that expanded clusters are preferentially positioned toward the synaptic center. To further confirm that these nanoscale changes were specific to trained animals, proximal-to-distal ratios of all the parameters were compared between control and trained animals (Figure 4G). In control animals, all metrics were centered around unity, indicating no intrinsic proximodistal difference. In contrast, trained animals exhibited significant differences in mean CPI (*p* = 0.03, n = 8 animals untrained, n = 6 trained animals across all comparisons, unpaired t-test), clustered fraction (*p* = 0.005), particles per cluster (*p* = 0.03), and cluster CPI (*p* = 0.02). Although some metrics did not reach statistical significance, overall results support a coordinated reorganization of GluA1 distribution following learning. We did not observe any similar intrasynaptic changes for GluA2 and GluA3 induced by the IA training (Supplementary Figures S8-9).

Together, these analyses demonstrate that learning promotes a nanoscale reorganization of GluA1-containing AMPARs characterized by moving toward the center and increased recruitment into expanded clustered nanodomains specifically in proximal CA1 synapses.

**Figure 4.**
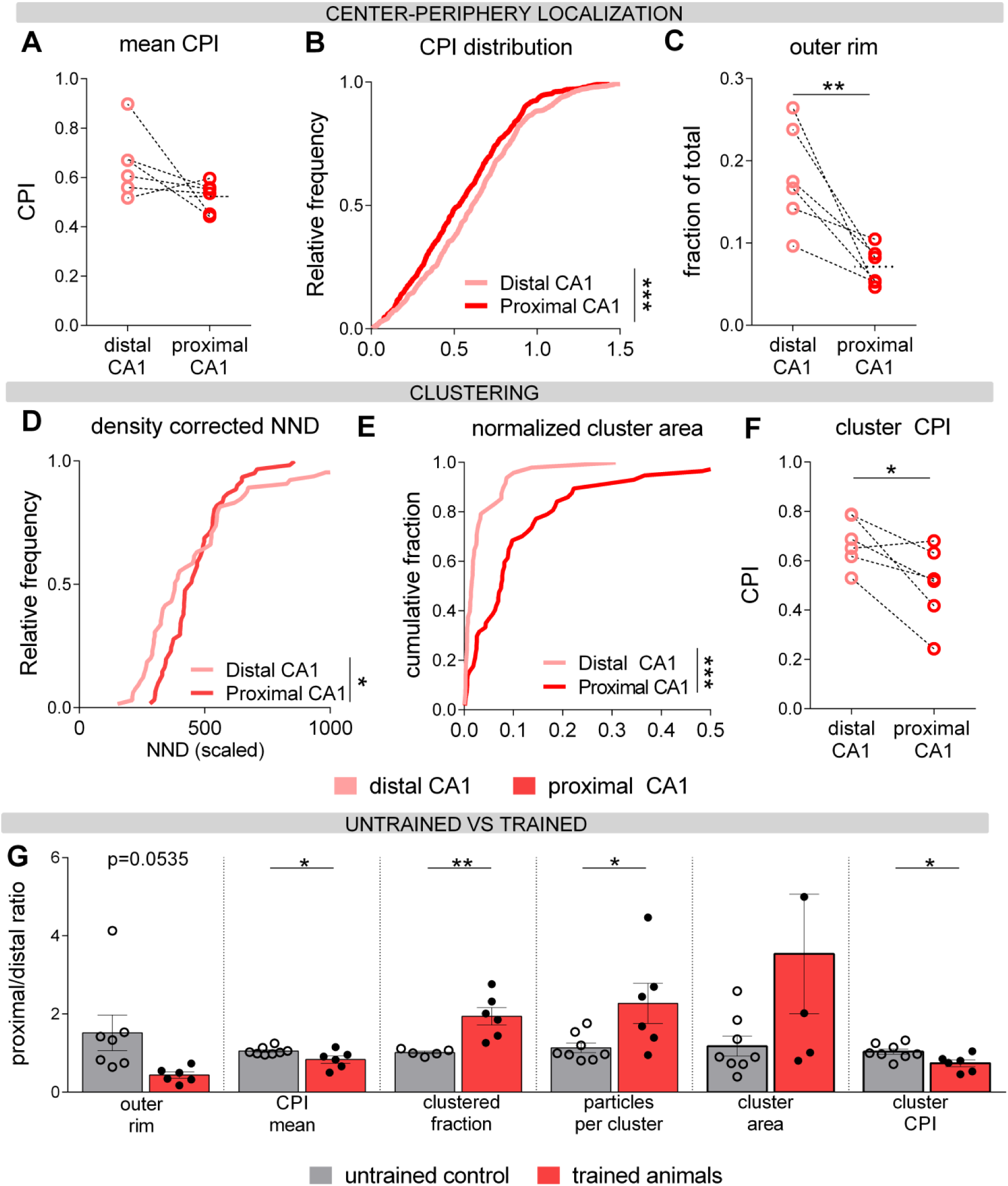
Learning drives nanoscale reorganization of GluA1 within synapses. **A.** Mean center–periphery index (CPI) values in proximal and distal synapses. **B.** Cumulative distributions of CPI values showing a shift toward more central localization in proximal synapses. **C.** Fraction of GluA1 particles located in the perisynaptic outer rim in proximal and distal CA1 synapses from trained animals. Points represent individual animals; lines connect paired measurements. **D.** Density-corrected nearest-neighbor distance (NND) cumulative distributions for proximal and distal CA1 synapses in trained animals. **E.** Density-corrected cluster area cumulative distributions showing a global shift toward larger clusters in proximal synapses. **F.** CPI of the cluster centroid indicates more central localization of the clusters in proximal than distal CA1. **G.** Proximal-to-distal ratios of nanoscale parameters in control and trained animals. Data are shown as individual animals (dots) with mean ± SEM. Statistical tests are described in the text.

## Discussion

Using SDS-FRL, the highest-resolution quantitative method available for *in situ* visualization of transmembrane proteins, combined with DAREA-based analysis, we demonstrate that inhibitory avoidance learning selectively increases synaptic GluA1 density and drives its redistribution toward the synaptic center in the proximal CA1 stratum radiatum. These findings provide, to our knowledge, the first direct ultrastructural evidence of intrasynaptic AMPAR reorganization following physiological, behavior-driven learning in the intact brain.

Our characterization of basal AMPAR distribution confirmed the highest labeling density in the stratum radiatum for all subunits, consistent with prior studies (46), while extending them to each GluA subunit at two-dimensional nanometer resolution. Notably, GluA3-containing receptors showed substantially lower density in extrasynaptic compartments — approximately 20–30% of GluA2 values — consistent with proteomic estimates that GluA3 represents around 20% of the total hippocampal AMPAR pool (47). This disproportionate enrichment of GluA3 at synapses relative to extrasynaptic membranes points toward distinct delivery mechanisms for GluA3-containing receptors, possibly related to the shorter intracellular C-tail of GluA3 and its differential interactions with synaptic scaffolding proteins compared to GluA1 (48,49).

At the intrasynaptic level, GluA1 and GluA2 labeling showed preferential peripheral localization, consistent with previous reports (50) while GluA3 was concentrated more centrally with fewer particles in the outer rim and greater clustering tendency, which we also reported in auditory relay synapses on bushy cells (33). This subunit-specific organization has direct functional implications. The peripheral enrichment of GluA1 and GluA2 is consistent with their proposed role in lateral diffusion-based synaptic incorporation during plasticity (26,51,52), whereas the central and clustered distribution of GluA3 — combined with its low extrasynaptic density — suggests a stably anchored pool more suited to sustaining basal transmission than participating in rapid potentiation (6). We did not observe discrete AMPAR nanocluster arrays despite strong clustering property for Cav2.1 in the presynaptic active zone in the CA1 area (30) under basal conditions, which is inconsistent with trans-synaptic nanocolumns (25). This may reflect genuine differences between perfusion fixed tissue and acute slice or cultured neuron preparations used in light microscopy studies, methodological differences in cluster detection, or the possibility that nanocolumn organization is a dynamic, activity-dependent phenomenon rather than a constitutive structural feature (27). It remains possible that GluA1-GluA2 heteromers are organized in nanocolumns but that this is difficult to resolve with GluA2 labeling, which also captures the centrally distributed GluA2-GluA3 population throughout the PSD.

Detecting molecular changes at individual synapses after physiological learning is inherently a search for a needle in a haystack. Only a sparse, distributed subset of CA1 synapses could be modified by a given experience (8,9), and by sampling randomly selected synapses per region in our measurements necessarily averages potentiated and naïve contacts — meaning the true magnitude of reorganization at genuinely modified synapses is certainly underestimated. That robust and statistically significant changes are nonetheless detectable, without genetic manipulation, pharmacological intervention, or artificial stimulation, attests to both the sensitivity of SDS-FRL and the coherence of the learning-induced effect. Within-animal normalization — proximal and distal measurements from the same replica samples — ensures that the observed proximodistal differences reflects genuine biological effects rather than technical artifacts.

The increased GluA1 density and concurrent trend toward decreased GluA2 density at synapses in the proximal CA1 area in trained animals, with no change in overall AMPAR content, is consistent with a shift in synaptic AMPAR composition from GluA1-GluA2 heteromers toward GluA1 homomers — receptors with higher single channel conductance (53,54) — in the initial phase of learning-induced plasticity. Such a compositional shift together with CaMKII-mediated GluA1 phosphorylation has been described following high-frequency electrical stimulation in acute slices and represents a current consensus view of early LTP expression (55,56), but has never been demonstrated at the ultrastructural level after physiological learning. This is consistent with the electrophysiological findings of Mitsushima et al. (22), where the increased AMPA/NMDA ratio was driven by increased AMPAR conductance rather than simply increased receptor number — a distinction our data now resolve at the molecular level.

Beyond changes in density and composition, intrasynaptic analysis revealed a shift in GluA1 localization toward the synaptic center in the proximal CA1 area of trained animals, detected as decreased center-periphery index and reduced outer rim fraction. This redistribution suggests that in the initial stages of learning-induced LTP, rearrangement of existing receptors toward the presynaptic release machinery may contribute to synaptic potentiation alongside the insertion of new GluA1 homomers, considering that receptors located close to the center of synapses would be exposed to higher glutamate concentration. More precise alignment of AMPARs with vesicle release sites enhances glutamate responses more effectively than simply increasing receptor number (23, 57), given the low glutamate affinity of AMPARs (58). Our data may extend this principle to GluA1-containing AMPARs in naturalistic memory formation in the intact brain. Taken together, our findings support a model in which learning-induced plasticity at CA3-CA1 synapses involves both a subunit composition change and a nanoscale spatial reorganization of the postsynaptic AMPARs — likely mediated by remodeling of the underlying scaffold through TARPs and PSD-95 (24,59,60) — rather than a simple increase in receptor number.

The present study has limitations. Critically, our sampling does not distinguish potentiated from naïve synapses. Future work combining inhibitory avoidance with activity-dependent synaptic tagging strategies (61,62) would enable selective ultrastructural analysis of learning-modified contacts — though current tagging approaches may require hours to days for reporter expression on the spine membrane detectable by SDS-FRL, and may not capture the rapid reorganization within 30 minutes of training described here. Developing faster activity-dependent labeling tools compatible with SDS-FRL will be essential for achieving true single-synapse resolution of plasticity in the intact brain. Nevertheless, our findings reposition synaptic nanoarchitecture, not receptor number alone, as one of the molecular substrates of memory.

## Data availability

The original electron microscopy images and associated quantitative data supporting the findings of this study will be deposited in a public repository. The accession number and the dataset will be made publicly available upon publication.

## Acknowledgments

This research received funding from the European Union (European Research Council Advanced grant no. 694539) and Human Brain Project SGA2 (Grant Agreement No. 720270) to R. S. The authors thank the staff of the Electron Microscopy Facility at Institute of Science and Technology Austria for their technical support and assistance during experimental work.

## Author contributions

M.J. and R.S. designed research; M.J., E.S., B.K., Y.S., L.S., and R.R. performed research; M.J., E.S., B.K., Y.S., L.S., and R.R. analyzed data; Y.F. and M.I. contributed new reagents/analytic tools; and M.J. and R.S. wrote the paper.

## Supplementary information

**Supplementary Figure S1.**
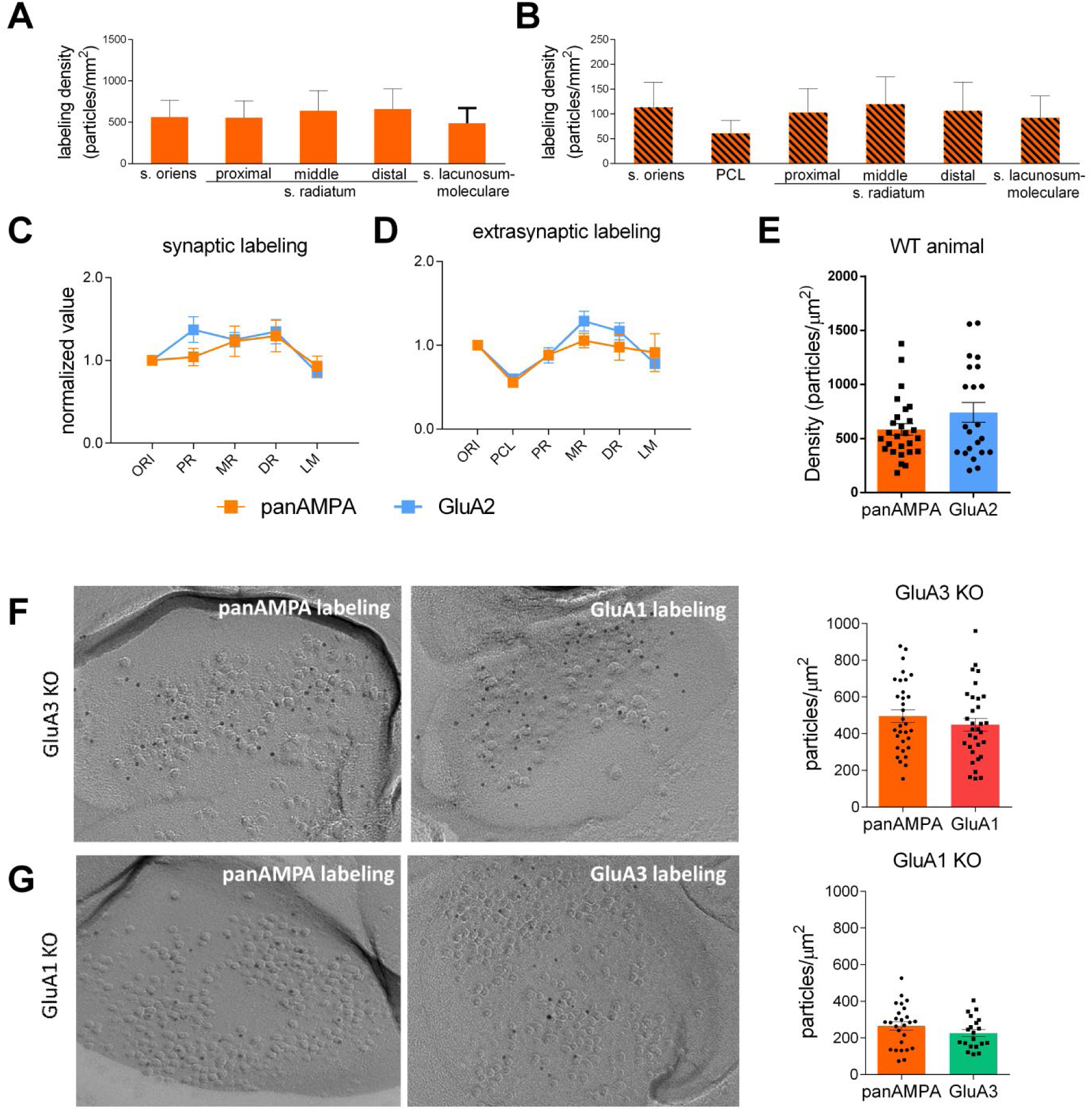
Labeling efficiencies of GluA subunit antibodies. **A-B**. Observed labeling densities for panAMPA (GluA1-3) in synaptic and extrasynaptic sites across CA1 layers. **C-D.** Comparison of normalized GluA2 and panAMPA labeling indicates similar distribution patterns across CA1 layers. **E-G.** Comparable labeling densities in complementary replicas prepared from WT, GluA3 KO, and, GluA1 KO animals indicate similar labeling efficiencies for panAMPA, GluA2, GluA1, and GluA3 antibodies (statistics described in the main text).

**Supplementary Figure 2.**
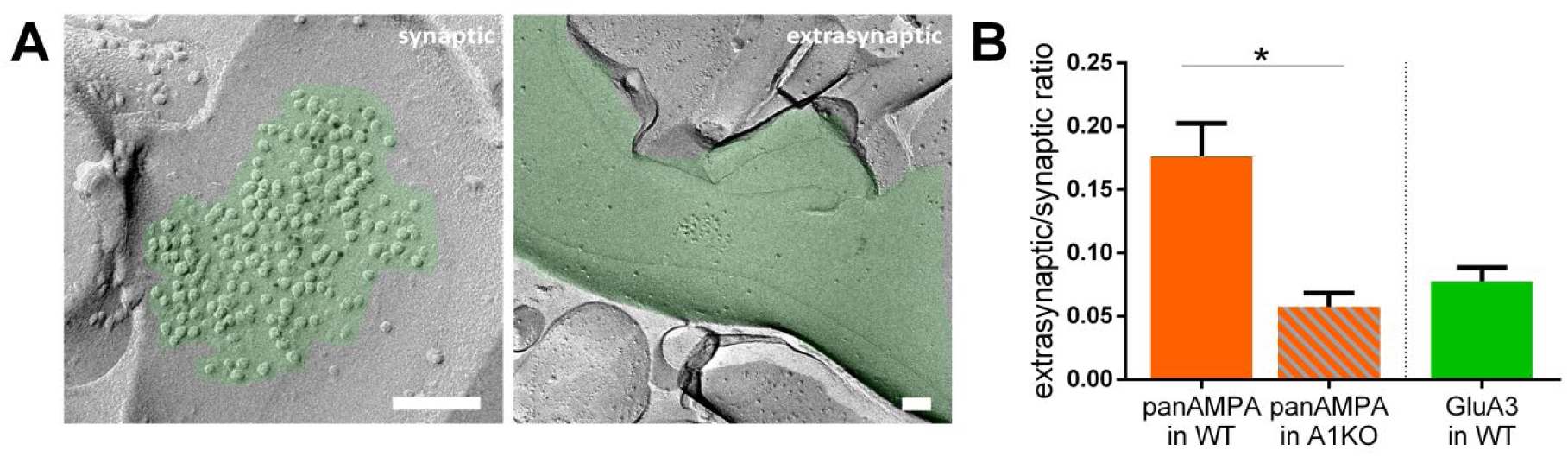
PanAMPA labeling in GluA1 KO animals. **A.** Representative images of panAMPA labeling in synaptic and extrasynaptic compartments show centrally localized labeling in synaptic sites and very little labeling in extrasynaptic sites. **B.** Significant reduction in extrasynaptic/synaptic ratios of panAMPA labeling in GluA1 KO compared with WT mice, and extrasynaptic/synaptic ratio of GluA3 in WT mice (the same data as shown in Figure 1G for comparison). Scale bars, 100 nm. Asterisks indicate levels of statistical significance: *p* < 0.05 (*), *p* < 0.01 (**), *p* < 0.001 (***).

**Supplementary Figure S3.**
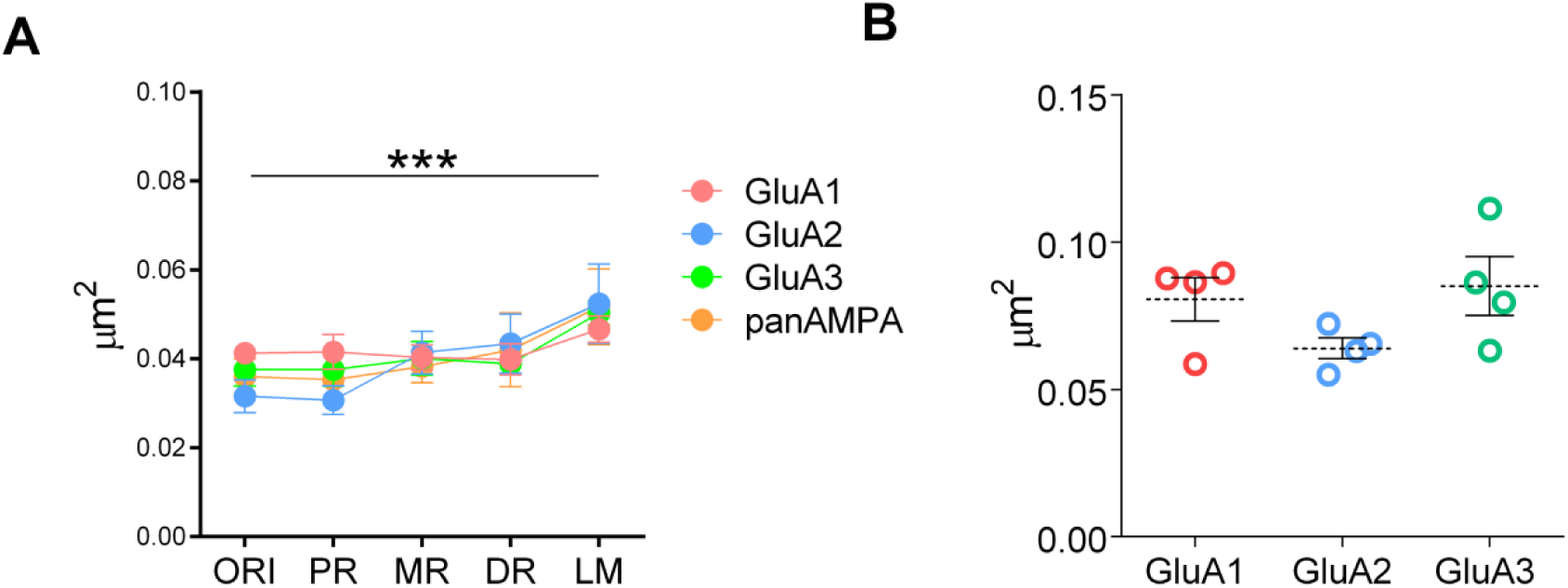
PSD areas across layers. **A.** No significant difference in the sampled PSD area across layers shows unbiased sampling of synapses for different subunits, while LM shows larger PSD areas than the other layers for all subunits. Corresponding to Figure 1. **B.** No significant difference in the sampled PSD area, corresponding to Figure 2.

**Supplementary table 1.** Post hoc comparisons related to Supplementary figure S3.

| Tukey's multiple comparisons test | Mean Diff, | 95% CI of diff, | Significant? | Summary | Adjusted P Value |
| --- | --- | --- | --- | --- | --- |
| ORI vs. PR | 0.0003100 | -0.009139 to 0.009759 | No | ns | > 0.9999 |
| ORI vs. MR | -0.0034 | -0.01285 to 0.006049 | No | ns | 0.8489 |
| ORI vs. DR | -0.004445 | -0.01389 to 0.005004 | No | ns | 0.6781 |
| ORI vs. LM | -0.01366 | -0.02311 to -0.004211 | Yes | ** | 0.0013 |
| PR vs. MR | -0.003710 | -0.01316 to 0.005739 | No | ns | 0.8036 |
| PR vs. DR | -0.004755 | -0.01420 to 0.004694 | No | ns | 0.6204 |
| PR vs. LM | -0.01397 | -0.02342 to -0.004521 | Yes | *** | 0.0010 |
| MR vs. DR | -0.001045 | -0.01049 to 0.008404 | No | ns | 0.9979 |
| MR vs. LM | -0.01026 | -0.01971 to -0.0008109 | Yes | * | 0.0268 |
| DR vs. LM | -0.009215 | -0.01866 to 0.0002341 | No | ns | 0.0593 |

**Supplementary table 2.** Post hoc comparisons related to Figure 1G.

| Tukey's multiple comparisons test | Mean Diff, | 95% CI of diff, | Significant? | Summary | Adjusted P Value |
| --- | --- | --- | --- | --- | --- |
| GluA1 vs. GluA2 | -0.008868 | -0.03933 to 0.02160 | No | ns | 0.7615 |
| GluA1 vs. GluA3 | 0.05480 | 0.02434 to 0.08527 | Yes | *** | 0.0002 |
| GluA2 vs. GluA3 | 0.06367 | 0.03320 to 0.09414 | Yes | **** | < 0.0001 |
| ORI vs. PR | 0.05809 | 0.01197 to 0.1042 | Yes | ** | 0.0071 |
| ORI vs. MR | 0.01250 | -0.03361 to 0.05862 | No | ns | 0.9378 |
| ORI vs. DR | 0.02152 | -0.02459 to 0.06763 | No | ns | 0.6767 |
| ORI vs. LM | 0.007253 | -0.03886 to 0.05337 | No | ns | 0.9915 |
| PR vs. MR | -0.04558 | -0.09170 to 0.0005286 | No | ns | 0.0541 |
| PR vs. DR | -0.03657 | -0.08268 to 0.009545 | No | ns | 0.1794 |
| PR vs. LM | -0.05083 | -0.09695 to -0.004721 | Yes | * | 0.0241 |
| MR vs. DR | 0.009017 | -0.03710 to 0.05513 | No | ns | 0.9807 |
| MR vs. LM | -0.005250 | -0.05136 to 0.04086 | No | ns | 0.9975 |
| DR vs. LM | -0.01427 | -0.06038 to 0.03185 | No | ns | 0.9031 |

**Supplementary Figure S4.**
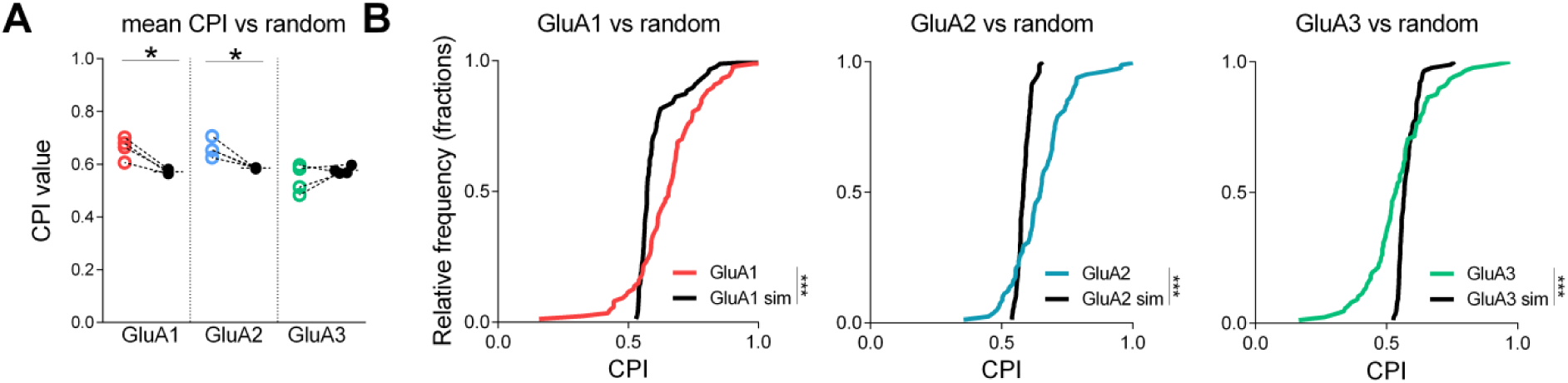
Comparison of real and simulated CPI values. **A.** Mean CPI comparison vs randomly distributed particles. GluA1 and GluA2 show significantly larger CPI values than those of randomly distributed particles. **B**. Cumulative CPI distributions for GluA subunits show significant differences from those for randomly distributed particles.

**Supplementary Figure S5.**
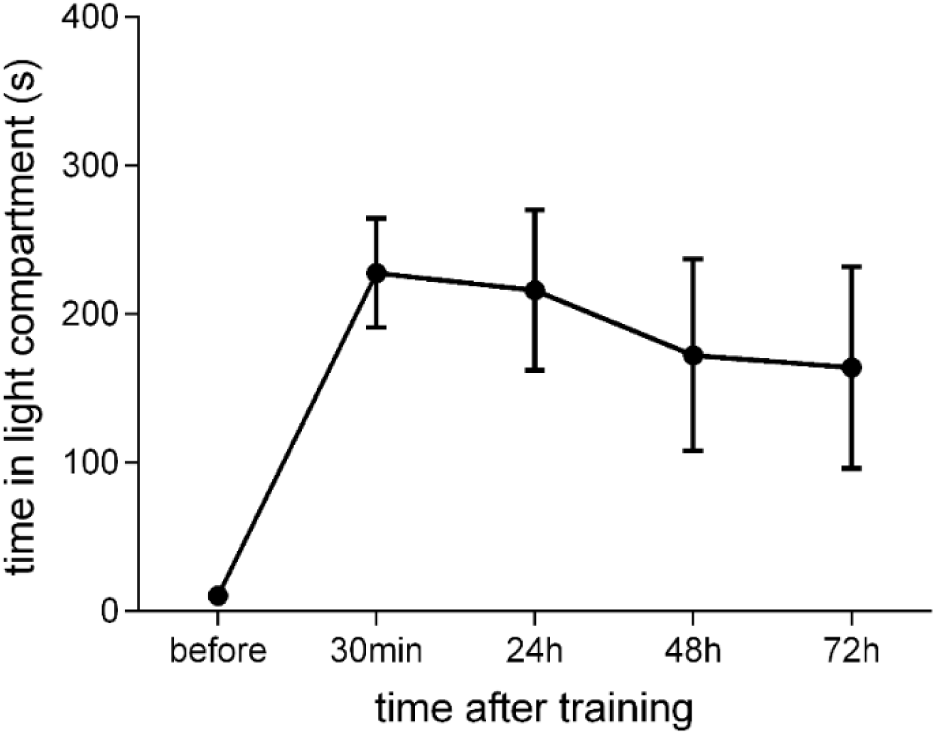
Inhibitory avoidance memory persisting over 72h.

**Supplementary Figure S6.**
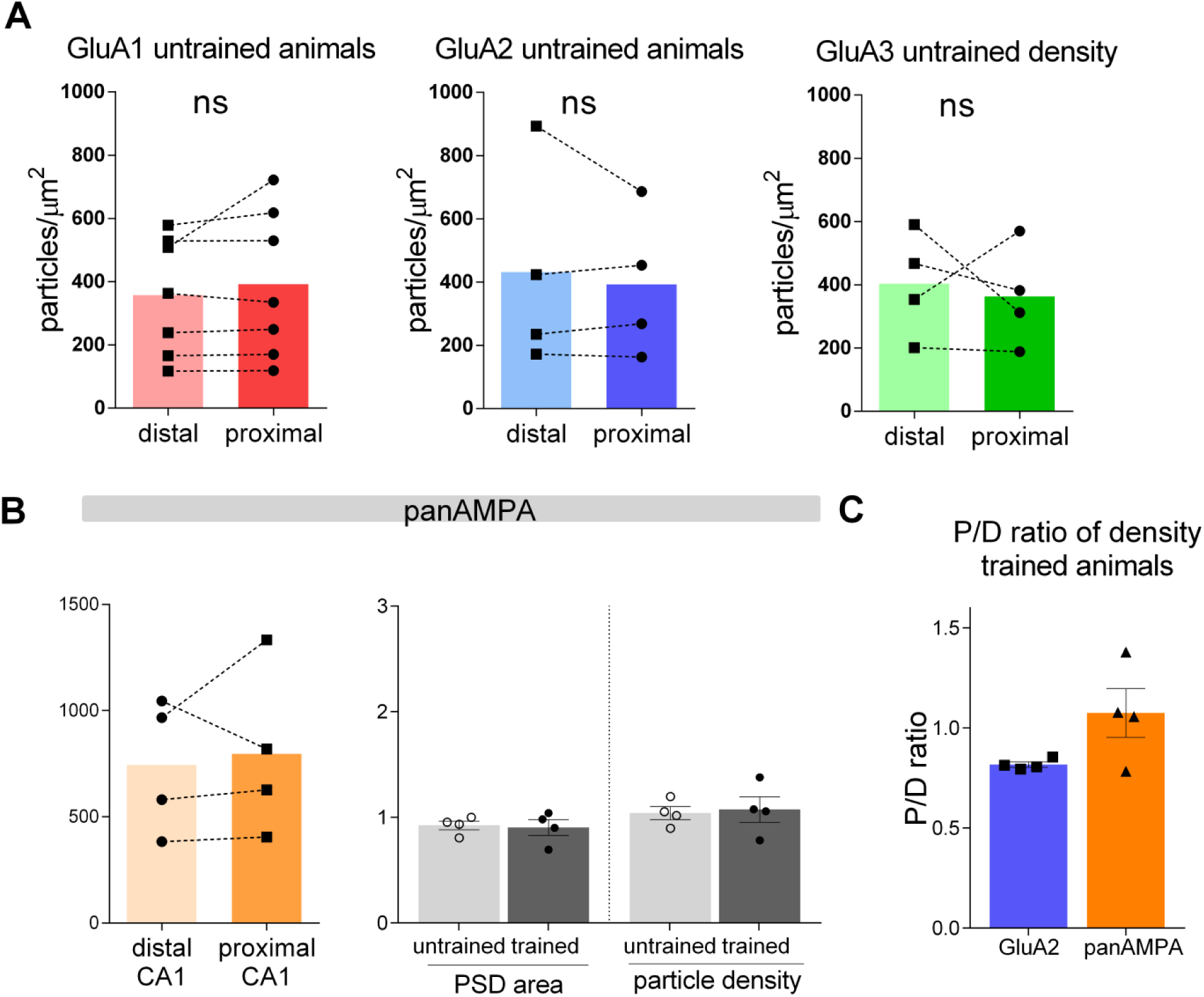
Labeling densities for GluA subunits in the proximal and distal CA areas. **A**. Comparison of P/D ratios of particle densities for GluA1-3 subunits in untrained animals. **B.** No significant difference in labeling densities for panAMPA and PSD areas between proximal and distal CA1 in untrained and trained animals. **C.** The P/D ratios of labeling densities in the CA1 area show a higher trend for panAMPA than GluA2 labeling (p = 0.54, unpaired t-test).

**Supplementary Figure S7.**
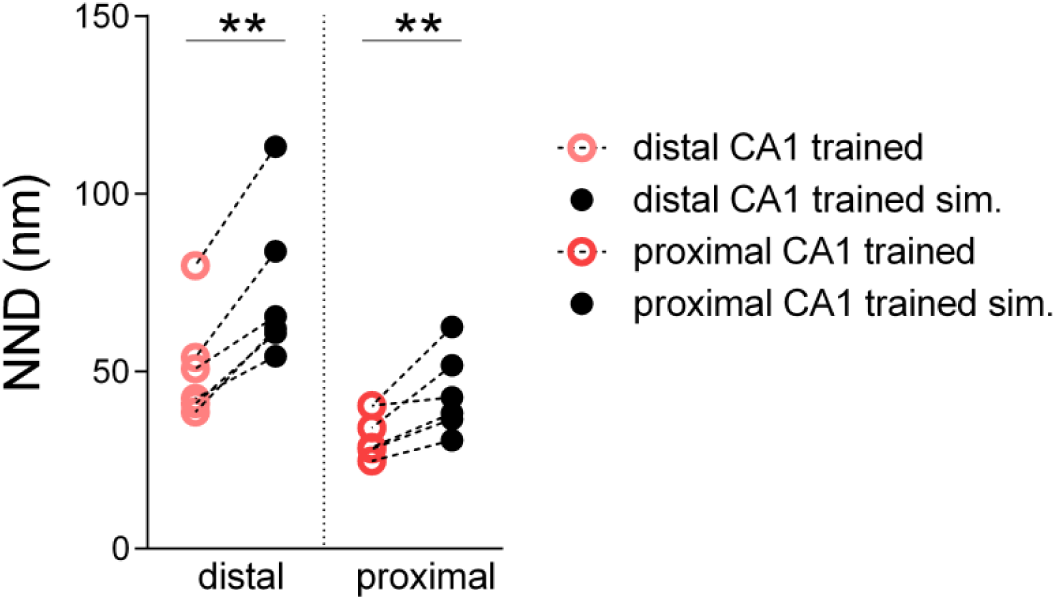
Comparison of NND values for real and simulated GluA1 particles in trained animals (distal vs sim, p = 0.001, proximal vs sim, p = 0.01, n = 6 animals for all groups, paired t-test).

**Supplementary Figure S8.**
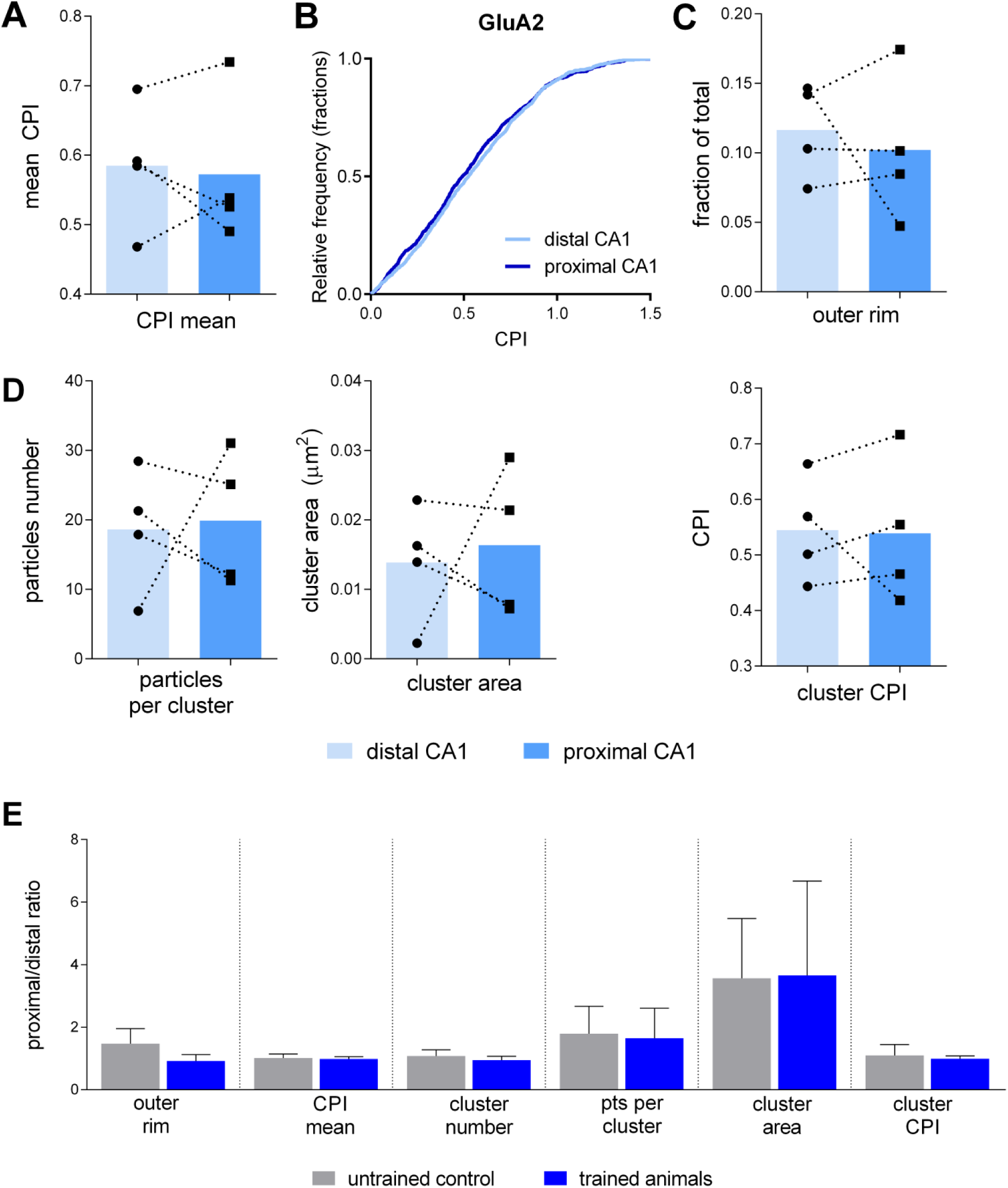
GluA2 distribution in IA trained animals. **A-D** Comparison of intrasynaptic parameters in trained animals. **E**. Comparison of proximal to distal ratios for various parameters between untrained and trained animals. No significant difference was detected between IA trained and untrained animals.

**Supplementary Figure S9.**
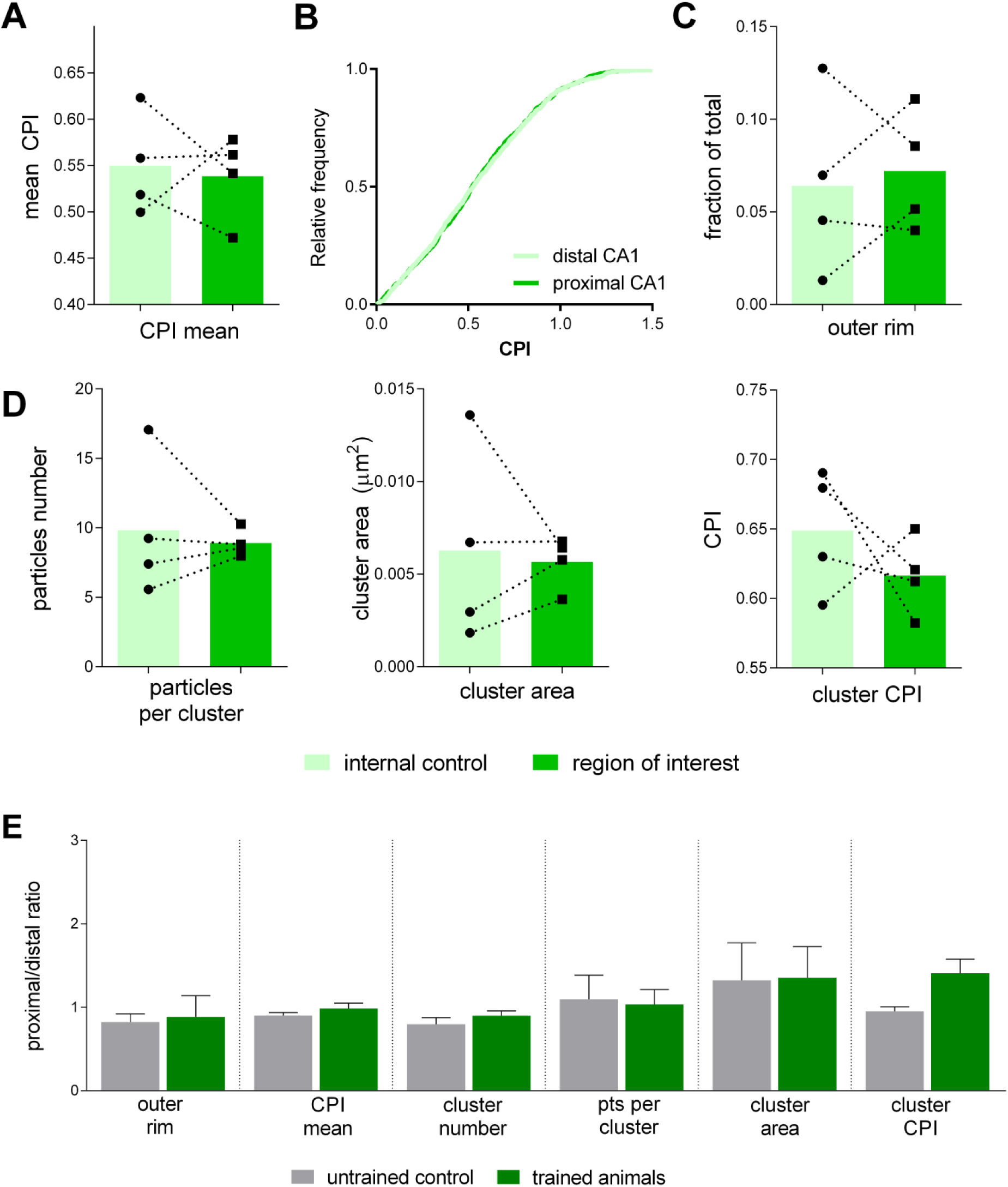
GluA3 distribution in IA trained animals. **A-D**. Comparison of intrasynaptic parameters in trained animals. **E.** Comparison of proximal to distal ratios for various parameters between untrained and trained animals. No significant difference was detected between IA trained and untrained animals.

